# Aberrant brain criticality as a neural basis of preclinical Alzheimer’s disease

**DOI:** 10.1101/2022.12.22.521549

**Authors:** Ehtasham Javed, Isabel Suárez-Méndez, Gianluca Susi, Juan Verdejo Román, J Matias Palva, Fernando Maestú, Satu Palva

## Abstract

Alzheimer’s disease (AD) is a chronic, nonlinearly progressive neurodegenerative disease that affects multiple domains of behaviour and is the most common form of dementia. However, there is scarce understanding of its biological basis nor there are reliable markers for the earliest disease stages preceding AD. Here we investigated whether AD progression is predicted by increasingly aberrant critical brain dynamics driven by underlying E/I imbalance using magnetoencephalography (MEG) data from cross-sectional (N=343) and longitudinal (N=45) cohorts. As a hallmark of brain criticality, we quantified long-range temporal correlations (LRTCs) in neuronal oscillations and tracked changes in neuronal excitability. We demonstrate that attenuation and progressive changes of LRTCs characterize the earliest stages of disease progression and yield accurate classification to individuals with subjective cognitive decline (SCD) and mild cognitive impairment (MCI). Our data indicate that pathological brain critical dynamics in AD progression provide a clinical marker for targeting specific treatments to individuals at increased risk.

## Main

Alzheimer’s disease (AD) is the most common form of dementia. It is a multifaceted disease characterized both by early memory loss but also various other forms of deficits in cognition, behaviour and social interactions (Hampel et al., 2022). It has been proposed that these cognitive impairments are partially attributable to alterations in large-scale network activity, as evidenced by stage-specific patterns of brain network dysfunction (Babiloni et al., 2016; Blinowska et al., 2017; López-Sanz et al., 2017). Functional hypersynchrony, in particular, has been found to be predictive of disease progression from MCI to AD (Maestú et al., 2021; Pusil et al., 2019) and also characterizes preclinical stages, such as in healthy relatives of AD patients (Ramírez-Toraño et al., 2021) and in subjects with high amyloid beta (Aβ) deposition (Nakamura et al., 2017). Correspondingly, hypersynchrony, a putative biomarker for AD, has been associated with Aβ deposition (Nakamura et al., 2017; Ranasinghe et al., 2022).

Such signatures of network dysfunction across the AD continuum could reflect underlying progressive changes in the neuronal excitation-inhibition (E/I) balance (Maestú et al., 2021) and alterations in the E/I ratio have indeed been proposed to be a key driving factor for brain network dysfunction in AD (Busche & Konnerth, 2016; Nifterick et al., 2023). In particular, dysregulation of the interplay between excitatory and inhibitory circuits may be associated with emerging neuronal hyper-excitability, putatively driven by Aβ deposition, which induces neurotoxicity at GABAergic terminals (Garcia-Marin et al., 2009). The E/I imbalance has been proposed as a mechanism for network dysfunction and cognitive impairment, and in animal models, therapeutic strategies that restore E/I balance have shown good promise to counteract neuronal dysfunction and cell loss (Busche & Konnerth, 2016). However, in humans, there is so far no direct electrophysiological evidence of altered E/I balance. Hypersynchrony *per se* (Pusil et al., 2019) cannot directly nor unambiguously be associated with hyper-excitability, as has been demonstrated with computational modeling (de Haan et al., 2017). However, while in the late stages of AD, subclinical or clinical epileptic activity is commonly observed, which can be seen as an indication of hyper-excitability, there is little neurophysiological evidence for such drastic E/I imbalance in the preclinical and prodromal stages.

Here, we tested the hypothesis that pathophysiology in AD and disease progression would be due to altered brain criticality throughout the disease continuum. Healthy brain activity is thought to have an operating point at or close to the critical transition between subcritical and supercritical phases in the system’s *state spac*e (Chialvo, 2010; Cocchi et al., 2017; Levina et al., 2009) i.e., between strong attenuation and strong amplification of activity propagation, with moderate levels of synchronization that are neither inadequate nor excessive (O’Byrne & Jerbi, 2022; J. M. Palva et al., 2018; Plenz & Thiagarajan, 2007). Operating at criticality endows a system with many functional benefits, such as maximal dynamic range, information transmission and representational capacity, all of which are instrumental to healthy brain function (Heiney et al., 2021; Zimmern, 2020). Brain criticality is assumed to be primarily controlled by the E/I ratio so that critical dynamics emerge at balanced E/I, while excess inhibition or excitation lead to sub- or supercritical dynamics, respectively (Plenz & Thiagarajan, 2007; Poil et al., 2012; Shew et al., 2009). In the super-critical regime, neuronal activity is overly amplified, allowing signals to propagate easily across the whole system, which is functionally detrimental (Amil & Verschure, 2021) and can lead to epileptic seizures (Gabriele Arnulfo et al., 2015; Meisel et al., 2015). Recent results converge on showing that healthy brains operating in the slightly subcritical phase very near the critical point (Toker et al., 2022; Wilting & Priesemann, 2019; Xu et al., 2022). We hypothesized that AD disease progression would be manifested in a progressive trajectory of the brain operating point (brain state) from the healthy state towards the supercritical regime due to excessive excitation.

Critical brain dynamics are evidenced by power-law long-range-temporal correlations (LRTCs) in oscillation amplitudes, i.e., power-law autocorrelations in amplitude fluctuations across lags of up to hundreds of seconds (Linkenkaer-Hansen et al., 2001; Nikulin et al., 2012; J. M. Palva et al., 2013; Poil et al., 2012; Simola et al., 2017; Smit et al., 2011; Zhigalov et al., 2015). We hypothesized that aberrant LRTCs and E/I imbalance could be potential novel mechanistic biomarkers for the putative earliest stages of AD, *i*.*e*., SCD and MCI. We obtained estimates of brain criticality (Linkenkaer-Hansen et al., 2001) and E/I balance (Bruining et al., 2020) for eyes-closed resting-state magnetoencephalography (MEG) recordings from neurotypical controls (NC) as well as SCD and MCI subjects. We first explored cohort-level differences in criticality and E/I metrics using cross-sectional data and then quantified their predictive value for the conversion from MCI to AD by using longitudinal data. Finally, to further assess the clinical potential of this approach, we used machine learning to classify early-stage AD patients employing brain criticality and E/I features.

## Results

### Demographics

The cross-sectional dataset included 116 neurotypical controls (NC), 85 participants with subjective cognitive decline (SCD), and 142 with mild cognitive impairment (MCI) (see Methods). Neuropsychological scores, general demographics, and differences between cohorts are presented in Table 1. The MCI participants (73.45±5.44 years) were slightly older than the SCD participants (72.16±5.29 years), who in turn were older than their NC peers (70.21±4.38 years). The age differences, albeit small, were significant between NC and SCD (*p*=0.013, independent samples t-test), and NC and MCI (*p*=0.003 ×10^−4^, independent samples t-test). There were no differences between cohorts in sex/gender. SCD and MCI participants performed significantly (p < 0.05, uncorrected) worse than NC in most neuropsychological tests (see Table 1). In each test, the performance of SCD participants lay within one standard deviation (SD) of the NC cohort and still within the range of performance considered ‘normal’. The MCI participants had much lower scores relative to the SCD and NC cohorts in all tests objectively demonstrating impaired performance in working memory, verbal fluency, and language function. When assessing medial temporal lobe (MTL) integrity, MCI participants showed lower volumes in all bilateral MTL regions (hippocampi, parahippocampi, and entorhinal cortices; normalized by the estimated total intracranial volume) than both SCD and NC participants (Extended data Fig. 1). Significant differences in left hippocampal and right parahippocampal volumes between NC (higher volumes) and SCD participants (lower volumes) were no longer significant after regressing out the age confound (Extended data Fig. 1).

**Table 1.**
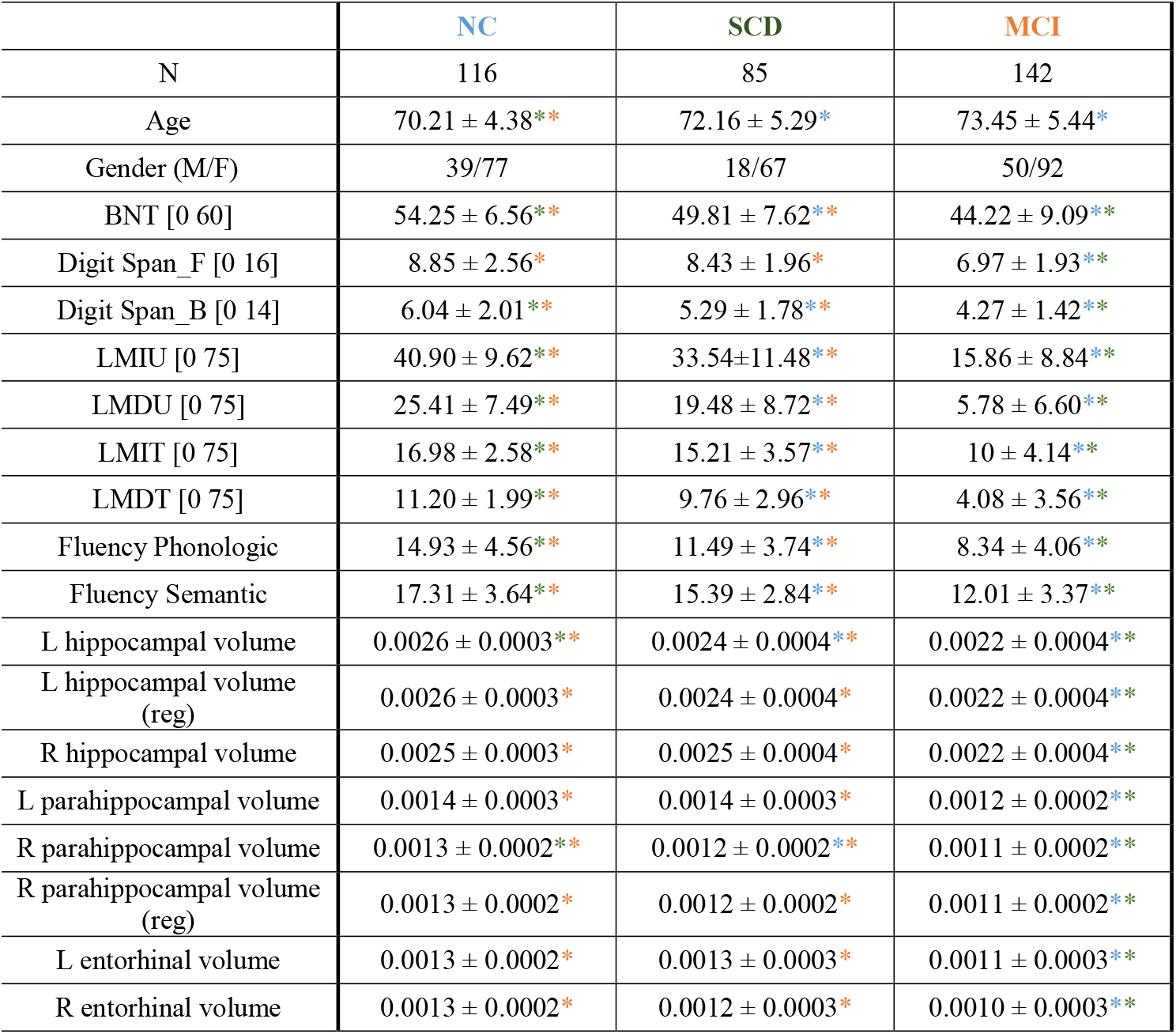
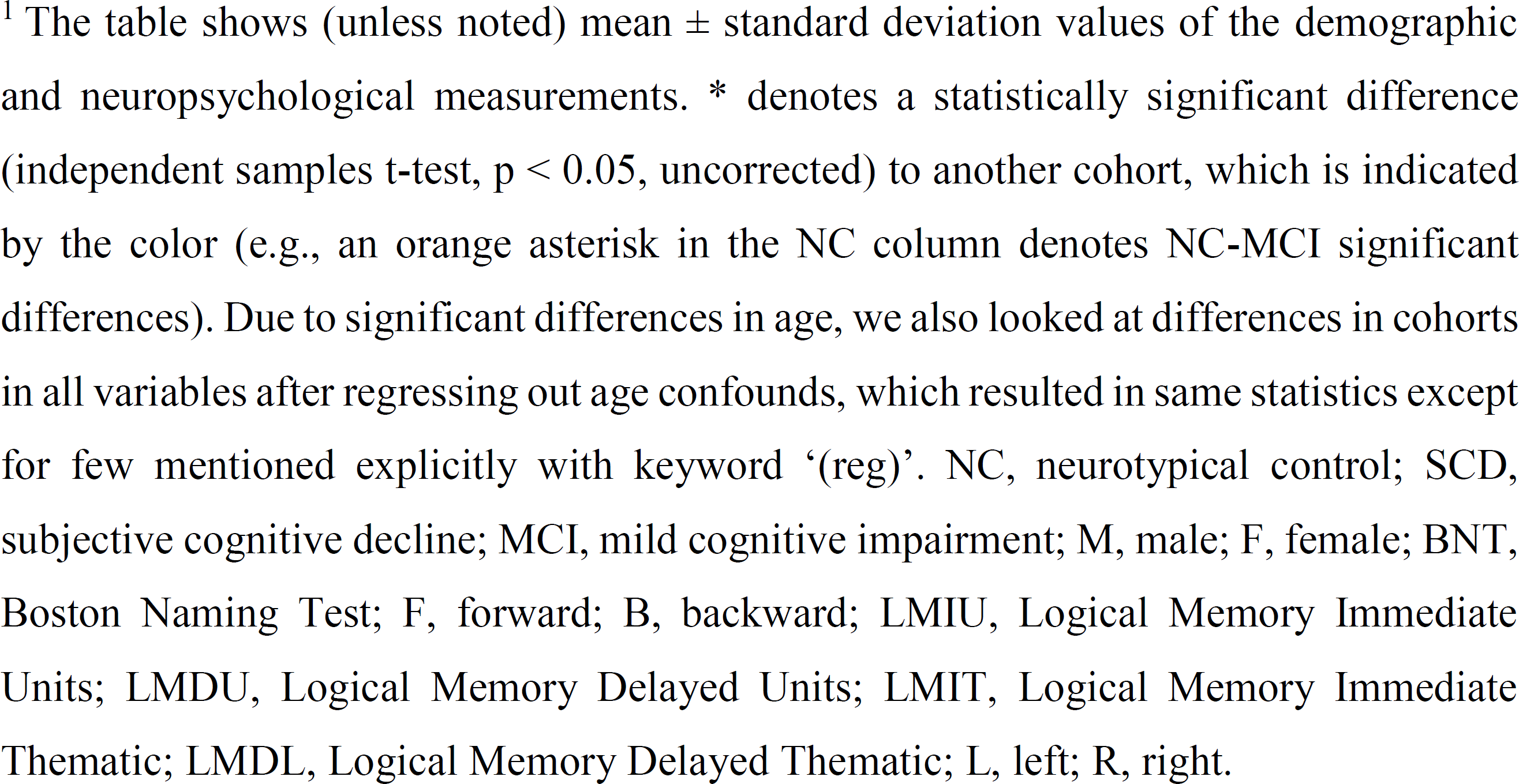
Demographic and neuropsychological measurements for each cohort in the cross-sectional dataset.

|  | NC | SCD | MCI |
| --- | --- | --- | --- |
| N | 116 | 85 | 142 |
| Age | 70.21 ± 4.38** | 72.16 ± 5.29* | 73.45 ± 5.44* |
| Gender (M/F) | 39/77 | 18/67 | 50/92 |
| BNT [0 60] | 54.25 ± 6.56** | 49.81 ± 7.62** | 44.22 ± 9.09** |
| Digit Span_F [0 16] | 8.85 ± 2.56* | 8.43 ± 1.96* | 6.97 ± 1.93** |
| Digit Span_B [0 14] | 6.04 ± 2.01** | 5.29 ± 1.78** | 4.27 ± 1.42** |
| LMIU [0 75] | 40.90 ± 9.62** | 33.54±11.48** | 15.86 ± 8.84** |
| LMDU [0 75] | 25.41 ± 7.49** | 19.48 ± 8.72** | 5.78 ± 6.60** |
| LMIT [0 75] | 16.98 ± 2.58** | 15.21 ± 3.57** | 10 ± 4.14** |
| LMDT [0 75] | 11.20 ± 1.99** | 9.76 ± 2.96** | 4.08 ± 3.56** |
| Fluency Phonologic | 14.93 ± 4.56** | 11.49 ± 3.74** | 8.34 ± 4.06** |
| Fluency Semantic | 17.31 ± 3.64** | 15.39 ± 2.84** | 12.01 ± 3.37** |
| L hippocampal volume | 0.0026 ± 0.0003** | 0.0024 ± 0.0004** | 0.0022 ± 0.0004** |
| L hippocampal volume (reg) | 0.0026 ± 0.0003* | 0.0024 ± 0.0004* | 0.0022 ± 0.0004** |
| R hippocampal volume | 0.0025 ± 0.0003* | 0.0025 ± 0.0004* | 0.0022 ± 0.0004** |
| L parahippocampal volume | 0.0014 ± 0.0003* | 0.0014 ± 0.0003* | 0.0012 ± 0.0002** |
| R parahippocampal volume | 0.0013 ± 0.0002** | 0.0012 ± 0.0002** | 0.0011 ± 0.0002** |
| R parahippocampal volume (reg) | 0.0013 ± 0.0002* | 0.0012 ± 0.0002* | 0.0011 ± 0.0002** |
| L entorhinal volume | 0.0013 ± 0.0002* | 0.0013 ± 0.0003* | 0.0011 ± 0.0003** |
| R entorhinal volume | 0.0013 ± 0.0002* | 0.0012 ± 0.0003* | 0.0010 ± 0.0003** |

<sup>1</sup> The table shows (unless noted) mean ± standard deviation values of the demographic and neuropsychological measurements. \* denotes a statistically significant difference (independent samples t-test, $p < 0.05$ , uncorrected) to another cohort, which is indicated by the color (e.g., an orange asterisk in the NC column denotes NC-MCI significant differences). Due to significant differences in age, we also looked at differences in cohorts in all variables after regressing out age confounds, which resulted in same statistics except for few mentioned explicitly with keyword '(reg)'. NC, neurotypical control; SCD, subjective cognitive decline; MCI, mild cognitive impairment; M, male; F, female; BNT, Boston Naming Test; F, forward; B, backward; LMIU, Logical Memory Immediate Units; LMDU, Logical Memory Delayed Units; LMIT, Logical Memory Immediate Thematic; LMDL, Logical Memory Delayed Thematic; L, left; R, right.

The longitudinal dataset comprised 45 participants that were followed up every 6 months (see Methods). At the end, 22 whose disease had not progressed to AD were diagnosed with stable MCI (sMCI) and 23 whose disease progressed to AD with progressive MCI (pMCI) based on clinical assessments. General demographics and Mini-Mental State Examination (MMSE) scores are presented in Extended data Table 1. There were no differences in sex/gender. At baseline, there were no significant differences between cohorts in age (sMCI: 71.73±5.20; pMCI: 73.70±3.40; *p*=0.138, independent samples t-test) or MMSE score (sMCI: 27.45±2.28; pMCI: 25.79±3.34; *p* = 0.067, independent samples t-test). In the post-session (the time between recordings: 1.98±0.85 years for pMCI and 2.33±0.57 years for sMCI cohort), there were significant differences between cohorts in age (sMCI: 73.05±4.85; pMCI: 75.87±3.55; *p*=0.032, independent samples t-test), which was expected since the duration of the follow-up depended on individual progression. Moreover, pMCI participants attained significantly worse post-session MMSE scores compared to sMCI participants (sMCI: 26.73±3.06; pMCI: 23.41±4.24; *p*=0.005, independent samples t-test) and to their baseline performance (pMCI (pre-session): 25.79±3.34; pMCI (post-session): 23.41±4.24; *p*=0.0008, paired t-test). Conversely, there were no significant differences on sMCI participants’ MMSE scores across sessions (sMCI (pre-session): 27.45±2.28; sMCI (post-session): 26.73±3.06; *p*=0.195, paired t-test).

### Attenuation of LRTCs dissociates early SCD and MCI stages

As an index of brain critical dynamics, we quantifed LRTCs in oscillation amplitude fluctuations using detrended fluctuation analysis (DFA) (Hardstone et al., 2012). DFA exponents were estimated for each participant and for all cortical parcels from parcel time-series for wavelet frequencies between 2−90 Hz (Fig. 1a; see Methods). Mean DFA exponents were then obtained by averaging the individual DFA exponents across all parcels and participants separately for the NC, SCD, and MCI cohorts. Mean DFA exponents showed a broad peak from alpha to beta frequencies (7−30 Hz) (Fig. 1b). Both SCD and MCI participants showed lower DFA exponents relative to the NC cohort from alpha to low-gamma frequencies (7−40 Hz) (Fig. 1b, Kruskal-Wallis test, *p*<0.05; False Discovery Rate (FDR) corrected across frequencies). Additional *post hoc* tests revealed a salient progressive attenuation of LRTCs with disease development, with significant differences in DFA exponents between the NC and SCD cohorts (hereafter, NC-SCD) in the alpha-band (7−12 Hz), and between the SCD and MCI cohorts (SCD-MCI) in the beta-band (12−22 Hz) (Fig. 1c-d, Fig. S1).

**Fig. 1.**
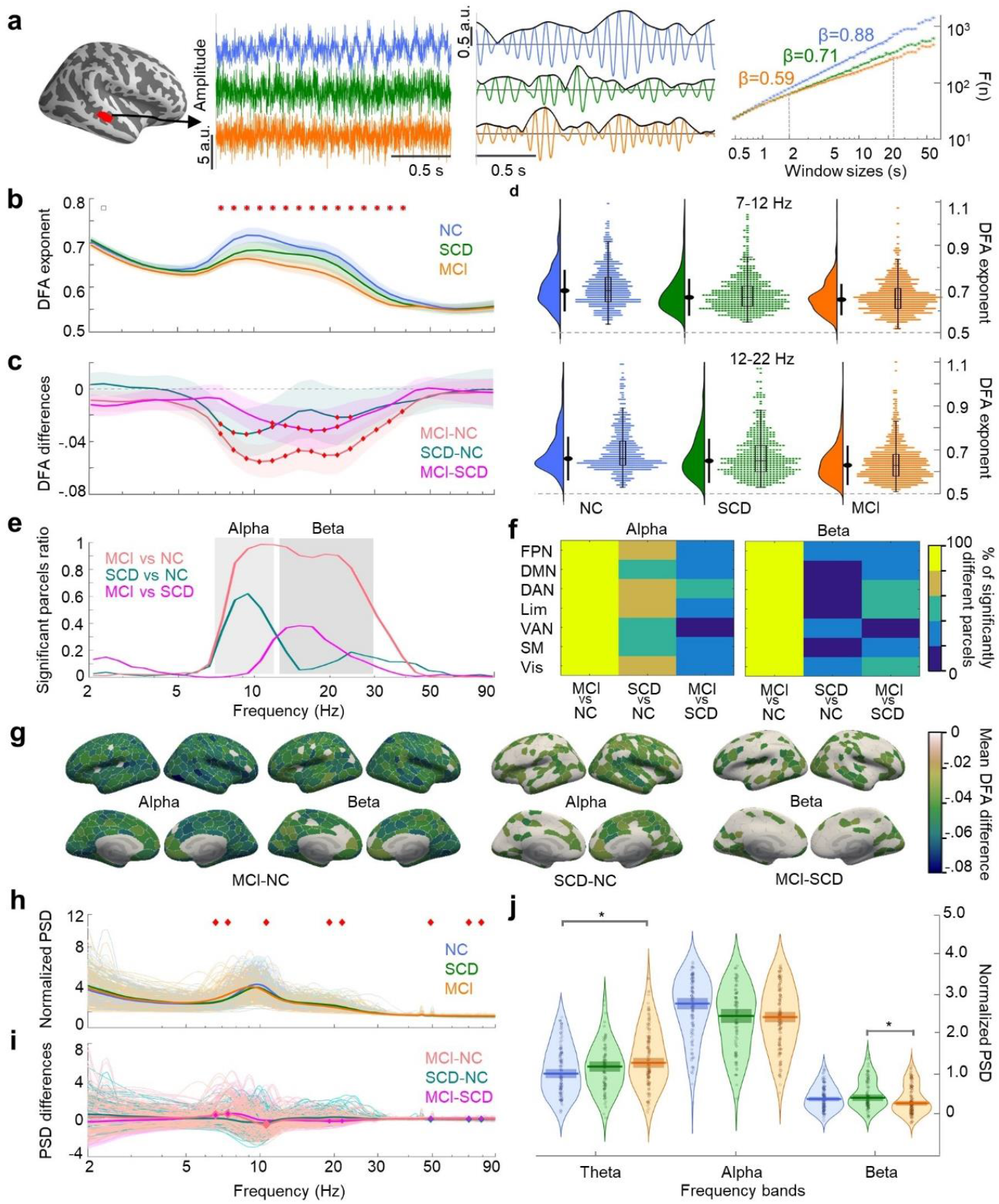
LRTCs dissociate NC, SCD, and MCI cohorts. **a**, Left: MEG broadband time-series from the superior temporal cortex (red parcel) for representative participants of each cohort (blue = Neurotypical controls (NC), green = Subjective Cognitive Decline (SCD), and orange = Mild Cognitive Impairment (MCI)). Middle: Narrow-band (10.6 Hz) parcel time-series with their amplitude envelopes. Right: Corresponding mean DFA exponents). **b**, Mean DFA exponent, averaged across parcels and within cohorts. Shaded areas represent bootstrapped (n=10,000) 95% confidence intervals. Red diamonds highlight the frequencies with significant differences (Kruskal-Wallis test, p<0.05, FDR corrected). **c**, Pairwise differences between cohorts in averaged DFA exponents. Red diamonds highlight significant differences. **d**, Density plots (left) for DFA exponents averaged within alpha (7□12 Hz) and beta (12□22 Hz) bands where SCD-NC and SCD-MCI differences were found within cohorts, respectively. The black-filled dot denotes median, and the line length represents the standard deviation. Individual participants’ DFA exponents (right) with an overlaid boxplot denoting the first, median and third quartiles, and the whisker lengths representing 1.5 times the interquartile range. **e**, Percentage of parcels showing statistically significant differences (Wilcoxon rank-sum test, *p*<0.05, FDR (Q=20%) corrected) in DFA exponents between NC-SCD, NC-MCI, and SCD-MCI. **f**, Differences between cohorts at the brain functional networks in the alpha and beta bands. **g**, Parcels that show pairwise differences for NC-MCI (left), NC-SCD (middle), and SCD-MCI (right). **h**, Normalized power-spectral-density (PSD), averaged across parcels and within cohorts. Red diamonds highlight differences between cohorts (Kruskal-Wallis test *p*<0.05, FDR corrected). **i**, Pairwise differences between cohorts in PSD (Wilcoxon-rank-sum test, FDR corrected). **j**, Mean PSDs averaged across theta, alpha, and beta bands with median (horizontal line) and 95% confidence intervals (shaded rectangle) for the assessment of band-wise differences. * denotes statistically significant differences (Wilcoxon-rank-sum test, *p<0*.*05*, Bonferroni corrected).

Taken that (i) AD is associated with structural changes that initially only take place in a subset of brain regions (Jack et al., 2013) and that (ii) DFA exponents appear to provide a robust measure of differences between cohorts in early-stage AD, we next examined the regional specificity by testing differences in DFA exponents at the parcel level. In line with the whole-brain results, differences in DFA exponents between the NC and MCI cohorts (hereafter, NC-MCI) were extensive and found in nearly all parcels in the alpha and beta bands (Fig. 1e). The NC-SCD and SCD-MCI differences showed expansion of pathological dynamics across disease progression, such that the early NC-SCD deterioration in DFA exponents occurred only in the alpha-band, while the progressing SCD-MCI differences appeared in the beta band, together overlapping with the observed NC-MCI differences (Fig. 1e). Region-wise NC-MCI differences were widespread and observed for all the brain functional systems (Fig. 1f) and parcels (Fig. 1g, Fig. S2), while NC-SCD and SCD-MCI differences showed regional specificity: NC-SCD alpha-band differences were visible in the frontoparietal network (FPN), dorsal attentional network (DAN), limbic network (Lim), and visual system (Vis), whereas SCD-MCI beta-band differences were localized within the DAN, Lim, and Vis networks (Fig. 1f-g).

### Elevated excitation characterizes disease progression

Theoretical models of critical brain dynamics suggest that scale-free LRTCs peak at around the critical point which represents balanced E/I. Excessive inhibition leads to a shift of the operating point towards the subcritical side and excessive excitation to the supercritical side of the critical point (Fig. 2a). To assess E/I balance, we used a recently developed method of *functional* E/I (*f*E/I) (Bruining et al., 2020), which estimates the E/I ratio from the relationship between LRTCs and the spectral power of neuronal oscillations (Fig. S3a; see Methods). The averaged fE/I values for each cohort differed between cohorts in alpha to gamma frequencies (Fig. 2b), Additional *post hoc* tests revealed that these differences were mainly due to a significant increase of *f*E/I in the MCI relative to the NC and SCD stages both at the whole brain (Fig. 2c) and at the parcel level (Fig. 2d). NC-MCI and SCD-MCI differences were spatio-spectrally pronounced while NC-SCD differences were overall weak (Fig. 2d), supporting the idea that E/I alterations echo disease progression and might represent a pathway of ongoing network disruption beginning at the SCD stage.

**Fig. 2.**
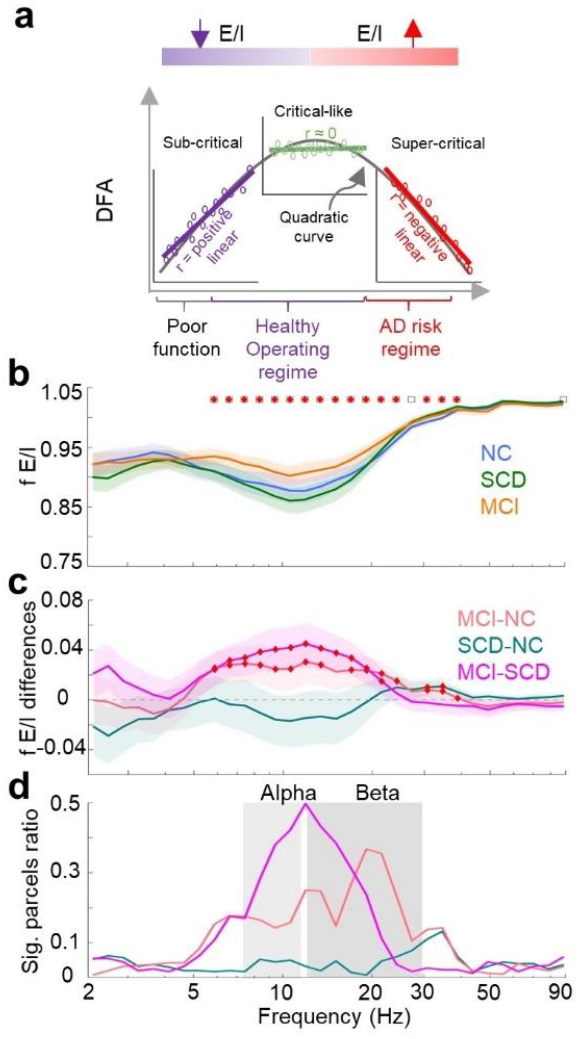
fE/I balance change across disease trajectory. **a**, Hypothesised dependence of DFA exponents on the brain critical state and the E/I balance in the classical brain criticality framework. Brain operating point is regulated by the E/I balance. DFA exponents peak at the critical transition point (inset: critical-like; green dots, no-linear dependence) at balanced E/I. The subcritical side (inset: purple dots, inhibition-dominant, positive-linear dependence) is characterized by stronger inhibition and the supercritical side (inset: red dots, excitation-dominant, negative-linear dependence) by stronger excitation. Quadratic dependence appears for all points across regimes **c**, Averaged mean fE/I values for each cohort (blue =NC, green = SCD, and orange = MCI) with shaded areas describing 95% confidence interval calculated using bootstrapping (n=10,000) method. Significance as in (Fig. 1b). **d**, Averaged pairwise differences between cohorts in the mean fE/I with 95% confidence intervals in bright and shaded colors, respectively. The diamonds mark the frequencies with significant (p<0.05; Kruskal-Wallis, FDR corrected) differences. **e**, Percentage of parcels showing significant differences between cohorts in fE/I. Shaded grey areas highlight frequencies in the alpha and beta bands.

To test this hypothesis in detail, we examined the covariance between DFA exponents and *f*E/I values (as in Fig. 2a), which is affected by the brain’s operating point in the critical regime (see also (Fuscà et al., 2022)). We found negative linear correlations (Fig. S3b, Pearson correlation coefficient, *p*<0.05; Bonferroni corrected) between DFA exponents and *f*E/I values for all cohorts in the alpha and beta band, in line with the DFA and fE/I alterations. The individuals in all cohorts were positioned along the fE/I axis from the subcritical to the supercritical side giving rise to a concurrence of linear negative and quadratic trend (Fig. S3c; right) at whole-brain level and at parcel level (Linear: Fig. S3d; left, and quadratic: Fig. S3d; right) as predicted by the hypothesis.

### Aberrant LRTCs are correlated with reduced MTL volumes and neuropsychological scores

The clinical diagnosis of probable AD dementia is generally achieved through validated neuropsychological tests, while biomarker evidence (e.g., disproportionate MTL atrophy) increases the certainty of issuing a reliable diagnosis (Hampel et al., 2022). Following this, we investigated whether our quantified aberrant brain dynamics were correlated with a decay in cognitive performance and MTL atrophy and could therefore be used as a biomarker. We restricted the analysis to the DFA exponents in alpha and beta-band frequencies where we had observed the largest and anatomically most widespread differences. We found a significant association between alpha-band DFA exponents and bilateral hippocampal (left: *r*=0.17, *p*=0.005; right: *r*=0.17, *p*=0.003), entorhinal (left: *r*=0.19, *p*=0.001; right: *r*=0.18, *p*=0.004), and parahippocampal volume (left: *r*=0.17, *p*=0.006; right: *r*=0.18, *p*=0.004) (Fig. 3a), and between beta-band DFA exponents and right entorhinal volume (*r*=0.16, *p*=0.004). The correlation of DFA exponents with neuropsychological scores (Table S1) was carried out at the parcel level. We found a positive correlation between alpha- and beta-band DFA exponents and memory performance (measured through nine neuropsychological scores) (Fig. 3b). These associations held true even after accounting for age (see Fig. S4). The topographical distribution of the correlations showed that in most parcels DFA exponents correlated with more than one neuropsychological score, revealing that these associations are not specific to any particular score (Fig. 3c).

**Fig. 3.**
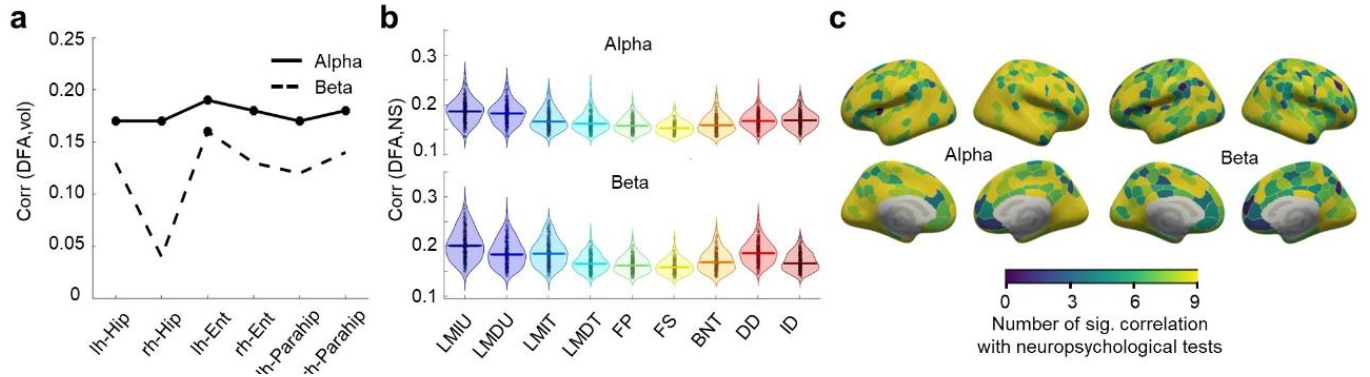
Aberrant LRTCs are correlated with structural and cognitive deficits. **a**, Correlation of DFA exponents with structural volume data (lh, left hemisphere; rh, right hemisphere; Hip, hippocampus; Parahip, parahippocampus; Ent, entorhinal cortex). **b**, Correlation between observed DFA exponents with neuropsychological scores as violin plots; the horizontal bar indicates the median. **c**, Cortical distribution of correlations with neuropsychological scores. A parcel’s color indicates the proportion of neuropsychological scores (out of 9) showing significant correlations with its DFA exponent.

### Classification of preclinical and prodromal AD

We showed that LRTCs and *f*E/I features provide a robust representation of differences between cohorts in early-stage AD. Next, we aimed to translate these differences into individual classifications for the different diagnostic categories of our study. This step is central for the discrimination of early-stage AD, since it has been shown that the traditional diagnosis based on neuropsychological scores could be challenged, e.g., in patients with high cognitive reserve (Lojo-Seoane et al., 2018). To this aim, k-nearest neighbors (kNN) classifiers were trained with minimal-optimal subsets of functional (LRTCs and *f*E/I) and structural (MTL volumes) features, selected by the mRMR algorithm (Fig. 4a, Fig. S5a; see Methods). MTL volume showed lower predictive power in the classification of SCD individuals compared to NC individuals, but became the most relevant feature in the prediction of MCI stage (Fig. 4b). Moreover, a subset of common features (e.g., fE/I values for SM in NC-MCI and NC-SCD; Fig.4b), inverted their tendencies along disease progression revealing a quadratic rather than monotonic path, similar to previous results (Pusil et al., 2019). Leave-one-out cross-validation yielded predictions of diagnostic categories greater than 80% (mean accuracy across all binary problems; (Fig. 4c)). Importantly, SCD individuals could be discriminated from both MCI and NC with high confidence (precision = 90% and 72%, recall = 80% and 84%, respectively). Similarly, the receiver-operating-characteristic (ROC) curves (Fig. 4d) endorsed the between-categories discriminative ability of the classifiers, with a mean area-under-curve (AUC) as high as 0.87. We also found that *f*E/I values were associated with higher predictive power during the early stage AD (NC-SCD), especially in theta- and gamma-bands, meaning that regionally specific differences are more vital for classifiers than widely-spread difference (alpha and beta bands). We further replicated the classifications by using support vector machine (SVM) method achieving similar classification performances (see Fig. S5b-d), demonstrating that results were robust against chosen methodology.

**Fig. 4.**
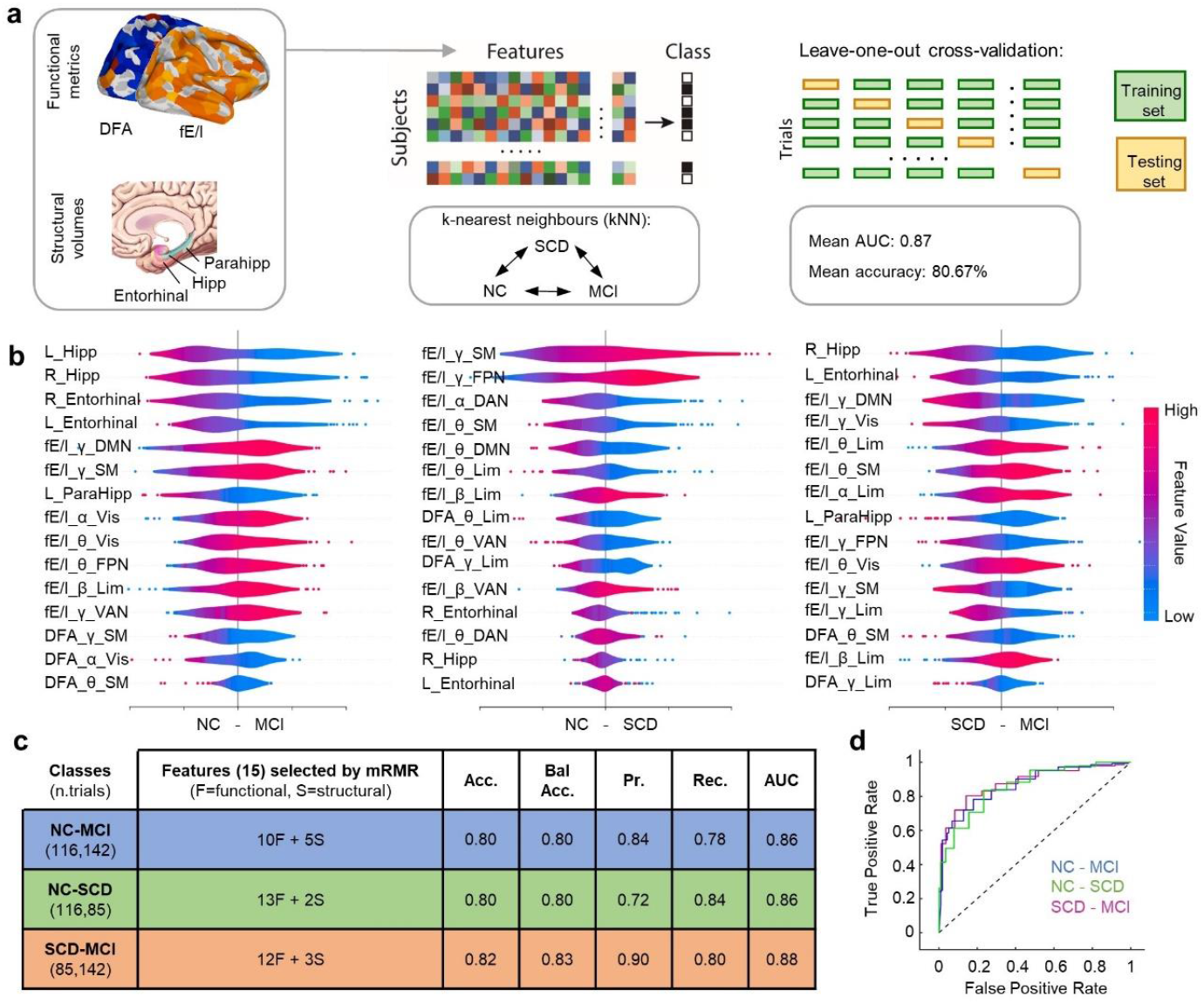
Preclinical and prodromal AD are efficiently discriminated based on neural data. **a**, An illustration of the functional (LRTCs and fE/I) and structural (MTL volumes) features employed by three separate kNN classifiers (k = 15) to classify pairs of the diagnostic categories included in the study (binary problems: NC-MCI, NC-SCD, and SCD-MCI). Classification was tested using leave-one-out cross-validation. **b**, SHAP summary plots for the different binary problems (NC-MCI, left; NC-SCD, middle; SCD-MCI, right) showing features ranked by importance in descending order and the SHAP values measure the impact of each feature on the model output (i.e., their predictive power). The color code shows whether high (red) or low (blue) values of the feature predict the target category. For example, in the NC-SCD summary plot, gamma-band fE/I in the sensorimotor network is the most predictive feature of the target category: SCD, and the color code shows that high values of that feature predict SCD. **c**, Summary of the classification results. **d**, ROC curve. Abbreviations: L, left; R, right; Hipp, hippocampus; ParaHipp, parahippocampus; Entorhinal, entorhinal cortex; θ, theta-band; α, alpha-band; β, beta-band; γ, gamma-band; DMN, default-mode network; SM, sensorimotor network; Vis, visual system; FPN, frontoparietal network; Lim, limbic network; VAN, ventral attentional network; DAN, dorsal attentional network; Acc, Accuracy, Bal Acc, balanced accuracy; Pr, precision; Rec, recall; AUC, area under curve.

### LRTCs and fE/I dissociate ‘stable’ and ‘progressive’ MCI patients

Finally, to test whether LRTCs and *f*E/I estimates would also predict disease progression within individuals, we analyzed a longitudinal dataset of MCI participants that were recorded twice (1^st^ and 2^nd^ measurements). After assessing their clinical status at a follow-up, MCI patients were divided into a *stable* (sMCI) group whose disease did not progress and a *progressive* (pMCI) group whose disease progressed to AD (see Methods). DFA exponents in frequencies 7−9 Hz and 30−35 Hz were significantly (p < 0.05, Kruskal-Wallis test) attenuated in the pMCI relative to the sMCI cohort already at the 1^st^ measurement during which the cognitive performance did not yet differ (Fig. 5a; upper left, see Extended data Fig. 2). Consistent with cross-sectional data results, DFA attenuations were spectrally extended (8-15 Hz) at the follow-up 2^nd^ measurement (Fig. 5a; upper-right). For sMCI, the DFA exponents remained stable from the 1^st^ to the 2^nd^ measurement and were significantly decreased only in the high-gamma band at the whole-brain level (Fig. 5a; bottom-left) and at the parcel level (Fig. 5b). In contrast, DFA exponents for the pMCI cohort were larger at 2^nd^ relative to the 1^st^ measurement for frequencies above 15 Hz both at the whole-brain level and at the parcel level. Differences in *f*E/I between cohorts at 1^st^ and 2^nd^ measurements, paralled those observed in DFA exponents, but however, did not differ yet at the 1^st^ measurement while in the 2^nd^ measurement fE/I values were larger for pMCI (Fig. 5c; upper right). In line, fE/I was changed in a wider frequency range between 1^st^ and 2^nd^ measurement for pMCI than sMCI cohort (Fig. 5c-d).

**Fig. 5.**
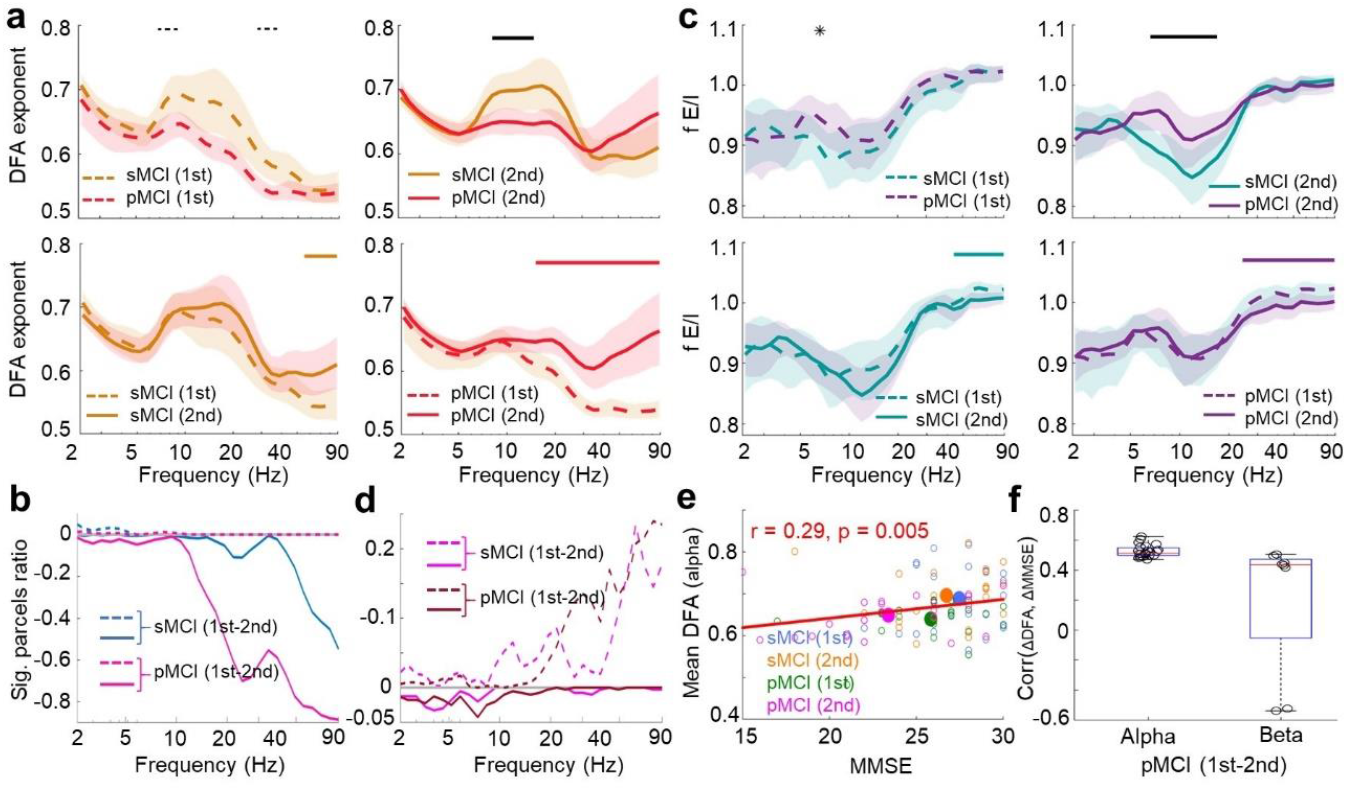
Longitudinal data: aberrant LRTCs and fE/I estimates predict disease progression. **a**, Mean DFA exponent differences between cohorts (stable (sMCI, N= 22) and progressive (pMCI, N = 23)) at 1^st^ (upper-left panel) and 2^nd^ (upper-right panel) measurements, and within-cohort differences between measurements for sMCI (bottom-left panel) and pMCI (bottom-right panel) with bootstrapped (n=10,000) 95% confidence intervals (shading). Statistically significant differences (Kruskal-Wallis test) between the cohorts are indicated with dotted black lines for 1^st^ measurement and solid black line for 2^nd^ measurement. Significant differences between 1^st^ and 2^nd^ measurements are indicated with brown and red lines for sMCI and pMCI cohorts, respectively. **b**, Proportion of parcels with a significant difference (1^st^ - 2^nd^) in measurements within cohorts (sMCI in blue and pMCI in magenta). Dotted line indicates the parcels that had DFA attenuated and solid lines the fraction of parcels with DFA increased, across measurements. **c**, The differences for fE/I, as in (a). **d**, The proportion of significant parcels for fE/I, as in (b). **e**, Correlation (Pearsons’ r) between participants’ mean alpha band DFA exponent and MMSE scores for both cohorts and both measurements. **f**, For pMCI, the change in MMSE between measurements is correlated with the change in DFA exponents in alpha and beta bands.

Importantly, DFA exponents (averaged across alpha band) across cohorts and measurements predicted the global cognitive status as measured with MMSE scores (Fig. 5e). Further, in alpha band, the differences in DFA exponents between 1^st^ and 2^nd^ measurement at the parcel level in the pMCI cohort predicted the decrease in MMSE scores with a positive linear correlation demonstrating that decreased DFA exponents were associated with decreased global cognitive status (Fig 5f) and thus demonstrated the functional significance of aberrant brain dynamics in predicting cognitive decline.

## Discussion

We addressed here a systems-level neurophysiological pathway leading to AD and hypothesized that aberrant LRTCs in oscillation amplitude fluctuations caused by E/I imbalance could characterize and enable the discrimination of early stages of AD (SCD and MCI). We analyzed LRTCs in the amplitude fluctuations of neuronal oscillations as an index of brain criticality and measure of emergent non-linear dynamics. We found that attenuated LRTCs in cortical oscillations characterize the early-stage AD and allow the dissociation of SCD and MCI individuals from healthy controls. Moreover, a progression of degraded LRTCs in alpha and beta frequencies discriminated SCD from MCI participants. Using longitudinal data, we further demonstrated that the conversion from MCI to AD could be predicted by these altered brain critical dynamics. In line with our recent hypothesis (Maestú et al., 2021; Pusil et al., 2019), stable and progressive MCI patients could be distinguished by the latter exhibiting a reduction in LRTCs paralleled with increased excitability and a shift from balanced or slightly inhibition-dominated (subcritical) brain dynamics towards excitation-dominated (supercritical) brain states and operating regimes. Our results are vital for understanding the AD pathophysiology with a focus on early diagnosis and intervention. In this respect, alterations in LRTCs in brain oscillations could serve as a novel biomarker for the early detection of individuals with SCD. Furthermore, since aberrant LRTCs also predicted conversion from MCI to AD, these measures could provide a clinical marker for targeting specific treatments to individuals at increased risk.

In light of the growing evidence of LRTCs being a proxy of brain criticality (Linkenkaer-Hansen et al., 2001; J. M. Palva et al., 2013; Poil et al., 2012; Zhigalov et al., 2015), our findings indicate that aberrant critical brain dynamics constitutes a key feature of early AD. More specifically, we show here that AD progression is associated with a shift of individual operating points towards the supercritical side. These findings have important implications for understanding of the neurophysiology of AD, as the primary control parameter regulating brain criticality is the E/I ratio (Plenz & Thiagarajan, 2007; Poil et al., 2012; Shew et al., 2009) and as LRTCs are influenced by neuromodulatory genes (Simola et al., 2017). Accordingly, healthy brain dynamics between insufficient and excessive neuronal synchronization emerge under conditions of balanced E/I (Chialvo, 2010; Levina et al., 2009; Poil et al., 2012), while disproportionate inhibition or excitation lead to sub- or supercritical dynamics, respectively (Amil & Verschure, 2021) and brain pathology (Babiloni et al., 2020; Busche & Konnerth, 2016; Meder et al., 2021). Healthy young adults have indeed been shown to operate in the slightly subcritical side of the critical point (Fuscà et al., 2022; Priesemann et al., 2014; Toker et al., 2022; Wilting & Priesemann, 2019). Here we show that healthy older adults, in contrast, operate around the critical transition point, suggesting that aging is associated with increased excitation. We further demonstrated that preclinical AD is associated with progressive transition towards the supercritical side of the critical transition point due to increased excitability compared to the NC cohort. This was further supported by the observed differences in LRTCs and fE/I estimates between the sMCI and pMCI patients.

In a recent framework, the “X model”, we proposed that alterations in the E/I balance lead to functional hyper-synchronization in prodromal AD patients (Pusil et al., 2019). This hypothesis was supported by data from animal models of the disease (Busche & Konnerth, 2016), showing that aberrant network interactions emerge as a consequence of exacerbated brain hyperexcitability, which might be partly due to the neurotoxic effects of Aβ plaques in the proximity of GABAergic synapses (Garcia-Marin et al., 2009). For the first time, we provide evidence for this hypothesis, showing that the SCD and MCI stages are associated with aberrant critical brain dynamics caused by increased hyperexcitability that grows throughout the disease continuum from sMCI to pMCI. However, the seemingly surprising longitudinal increase in LRTCs and decrease of fE/I in pMCI patients is in line with the non-linear U-shaped dynamics of brain criticality and the non-monotonic path as highlighted by the X-model, where an analogous trajectory was found for inter-areal phase synchronization (Pusil et al., 2019). These results suggest a severe pathological break-down of brain dynamics as the individual progresses towards AD. This is further corroborated by the observed correlation between change in measurements for LRTCs and MMSE scores.

As brain critical dynamics are correlated with synchronization of neuronal oscillations (Fuscà et al., 2022; S. Palva & Palva, 2012; Zhigalov et al., 2015), we propose that altered brain criticality leads to aberrant electrophysiological dynamics. Such alterations have been shown both in humans and animal models of AD (Babiloni et al., 2020; Maestú & Fernández, 2020) where AD patients typically exhibit a shift in the alpha peak frequency and a slowing down of rhythmic brain activity (Brassen & Adler, 2003; Jelic et al., 2000) and regionally and spectrally specific patterns of hyper- and hyposynchrony in both sensor-(López-Sanz et al., 2017; Maestú et al., 2015; Shumbayawonda et al., 2020) and source-level analysis (Babiloni et al., 2016; Blinowska et al., 2017). Although MCI patients showed aberrant LRTCs in a wide range of frequencies, in line with a previously observed frequency shift in AD (Brassen & Adler, 2003; Jelic et al., 2000), we found that in the cross-sectional analysis, LRTCs in alpha oscillations were attenuated for SCD, and upon disease progression to MCI, this attenuation was extended to beta oscillations. This extension supports the model of hyperexcitation, which has been reported in neurophysiologically-informed computational models, as increased excitation would lead to a shift in rhythmic activity from alpha to beta frequencies (Pinto et al., 2003). Given that neuronal oscillations regulate information processing and communication across distributed brain areas (G Arnulfo et al., 2020; Fries, 2015; Singer, 1999; Womelsdorf et al., 2014) serving fundamental functional roles in a variety of human sensory and cognitive functions (S. Palva & Palva, 2012; Siegel et al., 2012; Thut et al., 2012), aberrant brain criticality is likely to lead to cognitive deficits characteristic to early stage AD via its relationship with synchronization. In line with this notion, the strongest deficits in critical brain dynamics in SCD and MCI were observed in DMN and DAN, which coordinate attentional and executive functions (Power & Petersen, 2013), and exhibit both AD-related alterations in functional connectivity (López-Sanz et al., 2017; Pini, 2023) and accumulation of Aβ plaques on inhibitory terminals (Buckner et al., 2009).

Previous classification studies based on functional measures, such as reduced oscillatory power (Fernández et al., 2013) and functional connectivity (Maestú et al., 2015), have shown that MCI can be dissociated from AD with high accuracy of 79%–83% (Pusil et al., 2019), but distinguishing SCD from NC and/or MCI with such accuracies, is an ongoing quest. Our results, obtained by decoding with k-NN classifier using functional and structural neuroimaging data, demonstrated that both MCI and SCD patients can be accurately classified using non-linear brain dynamics measures while the more classical structural measures were more relevant for MCI but not the SCD patients. Our results demonstrate that critical brain dynamics may serve as a novel biomarker for early and prodromal stages of AD. These findings could open new avenues for pharmacological and other interventions targeted to recovery and sustainability of E/I balance, thereby preventing aberrant brain dynamics and excitotoxicity. Indeed, in animal models, the reduction of hyper-excitability by an antiepileptic drug has been shown to lead to improved cognitive abilities (Sanchez et al, 2008). Non-pharmacological approaches such as brain neuromodulation could base their hypothesis as well on the current findings, targeting specific frequencies and E/I profiles.

## Methods

### Study population

The study population included 116 NC (age: 70.21 ± 4.38 years (mean ± standard deviation); females: 66.38%), 85 SCD (age: 72.16 ± 5.29 years; females: 78.82%), and 142 MCI (age: 73.45 ± 5.44 years; females: 64.79%) participants (see Table 1). SCD diagnosis was determined using the SCD-I criteria: (i) self-reported cognitive concerns (mainly associated with memory); (ii) normal performance on standardized cognitive testing; (iii) feeling that the cognitive decline impairs everyday function; (iv) seeking of medical consultation; and (v) age of 60 years or more at the onset of SCD (occurring within the last 5 years). The cognitive decline was also confirmed by a reliable informant. Definitive inclusion in this cohort was decided by multidisciplinary experts after discarding potential confounders of SCD (e.g., psychiatric/neurological disorders, medical conditions, prescribed drugs, or substance use). MCI diagnosis was determined using the NIA-AA guidelines for MCI due to AD (with intermediate likelihood) (Albert et al., 2011): (i) self- or informant-reported cognitive concerns; (ii) objective evidence of impairment in one or more cognitive domains; (iii) preserved independence in everyday function; (iv) absence of dementia and (v) evidence of neuronal injury (hippocampal volume measured by MRI). Besides, additional clinical tests were conducted to discard other potential causes of cognitive decline (e.g., B12 vitamin deficiency, diabetes mellitus, thyroid disease, syphilis, and human immunodeficiency virus). Participants in the NC cohort were cognitively healthy and did not report cognitive complaints. General exclusion criteria also involved: (i) history of psychiatric/neurological disorders, or drug consumption that could impact MEG activity (e.g., cholinesterase inhibitors); (ii) evidence of infection, infarction, or focal lesions in a T2-weighted MRI scan within 2 months to the MEG recording (rated by two independent radiologists); (iii) modified Hachinski score ≥ 5; (iv) Geriatric Depression Scale-Short Form score ≥ 5; and (v) history of alcoholism/chronic use of anxiolytics, neuroleptics, narcotics, anticonvulsants, or sedative hypnotics. A neuropsychological battery was available for all participants including Boston Naming Test (BNT), Phonemic and Semantic Fluency Tests, Digit Span Test (forward and backward) and Texts of Verbal Memory of the Wechsler Memory Scale-III (Spanish version). To analyze AD conversion, 45 MCI participants were followed-up every 6 months and classified according to their clinical status as ‘progressive’ MCI (pMCI) (if meeting the criteria for probable AD (Pusil et al., 2019) at follow-up; 23 participants; female: 60.86%) or ‘stable’ MCI (sMCI) (if still meeting the criteria for MCI by the end of follow-up; 22 participants; female: 63.63%) (see Table 2). All the participants were recorded over 2 MEG measurements (1st- and 2nd-session) with an inter-session interval of 1.76 ± 0.70 for the sMCI cohort (age: 71.73 ± 5.20 years (pre-session); 73.05 ± 4.85 years (post-session)) and 2.17 ± 1.03 for the pMCI cohort (age: 73.70 ± 3.40 years (pre-session); 75.87 ± 3.56 years (post-session)). Pre- and post-session MMSE scores were recorded for every participant.

All the participants were right-handed and Caucasian Spanish speakers. All participants signed informed consent before enrolling. The study was approved by the Ethics Committee of the Hospital Clínico Universitario San Carlos and conducted according to current guidelines and regulations.

### Data acquisition

Eyes-closed resting-state MEG data were recorded using a 306-channel (102 magnetometers, 204 planar gradiometers) Vectorview system (Elekta AB, Stockholm, Sweden) installed at the Center for Biomedical Technology (Madrid, Spain). MEG data were sampled at Fs =1,000 Hz with an online anti-alias band-pass filter (0.1−330 Hz). The recordings lasted from 3 to 5 minutes. Additionally, three-dimensional T1-weighted anatomical MRI scans were acquired in a 1.5 T MRI system (GE Healthcare, Chicago, Illinois) using a high-resolution antenna and a homogenization PURE filter (fast spoiled gradient echo sequence, with parameters: repetition time/echo time/inversion time = 11.2/4.2/450 ms, flip angle = 12°, slice thickness = 1 mm, 256 × 256 matrix, and field of view = 256 mm). MRI data were collected at the Hospital Clínico Universitario San Carlos (Madrid, Spain).

### MEG data pre-processing

MEG data were pre-processed in two steps at two different sites, the Complutense University of Madrid (UCM) (segmented data pre-processing) and the University of Helsinki (UH) (continuous data pre-processing), using MATLAB and the Fieldtrip toolbox (https://www.fieldtriptoolbox.org/). Temporally extended signal space separation method (tSSS, Maxfilter software) was to remove external magnetic noises, interpolate bad channels, and compensate for head motion during the recording. Date were segmented into 4-s epochs of non-overlapping activity and segments containing muscular, ocular (blinks), or SQUID jump artifacts were annotated and independent component analysis (ICA) was applied to clean the data. Additional ‘high-amplitude’ artifacts were annotated and replaced with interpolated data.

### Source-reconstruction and collapsed parcel time-series

Source reconstruction was performed with minimum norm estimation (MNE) using dynamic statistical parametric maps (dSPM). The estimated dipole sources were then collapsed into Schaefer’s 400 parcels after applying a procedure of *fidelity weighting* to minimize the contribution of spurious inter-areal interactions (Siebenhühner et al., 2020). Data were then filtered with 32 Morlet wavelets (width parameter *m* = 5) at log-equidistant spacing within a range of 2−90 Hz.

### Analysis of DFA exponents

The DFA exponents were calculated as in (J. M. Palva et al., 2013; Zhigalov et al., 2015) for each wavelet frequency and parcel. The fitting interval included window sizes from 2−25 seconds (with 50% overlap) and robust bi-square fitting yielded the estimated scaling exponent.

### Analysis of functional excitation-inhibition ratio (fE/I)

Custom MATLAB scripts were implemented to compute dE/I as in (Bruining et al., 2020) for each parcel time-series: (i) parcel time-series were wavelet-filtered into 32 narrow-band signals as described above and their amplitude envelopes were extracted; (ii) the cumulative sum of each (demeaned) signal was calculated (the signal profile); (iii) the signal profile was divided by its mean amplitude in fixed windows of 40 cycles, to remove the effect of the original signal magnitude; (iv) each normalized signal profile window was linearly detrended; (v) the standard deviation was calculated for each window to get the windowed-normalized fluctuation function *w_nF*(*t*); (vi) mean windowed amplitudes (*wAmp*) were estimated (as in iii); (vii) the Pearson correlation between *w_nF*(*t*) and *wAmp* was calculated, that is, corr(*w_nF*(*t*), *wAmp*); and (viii) *f*E/I values were quantified as fE/I = 1 − corr(*w_nF*(*t*), *wAmp*), so that *f*E/I = 1 indicates balanced E/I, *f*E/I < 1 indicates dominant inhibition, and *f*E/I > 1 indicates dominant excitation.

### Classification model

kNN classification algorithm (*k* =15) was used to classify individuals based on a set of functional (LRTCs and *f*E/I) and structural (bilateral MTL volumes) features. Three independent classifiers were implemented based on the diagnostic cohorts in this study (i.e., NC vs. MCI, NC vs. SCD, and SCD vs. MCI). The initial set of features included: 56 functional features (i.e., FPN, DMN, DAN, Lim, VAN, SM, Vis), in theta (4-7 Hz), alpha (7-12 Hz), beta (12-30 Hz), and gamma (30-45 Hz) frequency bands; and 6 structural features (i.e., bilateral MTL volumes: hippocampi, parahippocampi, and entorhinal cortices) each example (trial) representing one participant (Fig.S4a). Data dimensionality was reduced by applying the *feature selection* algorithm (mRMR; (Radovic et al., 2017)) to select a minimal-optimal subset of 15 features for each of the three binary problems (see Fig. 4b). To counteract the negative effects of possible anomalous patterns on the learning procedure, the trials used for feature selection were previously passed through an outlier detection procedure, so that trials including values ≥ 3 of scaled median absolute deviations from the related feature’s median were temporarily discarded. Once the features were selected, rejected trials were re-introduced in the dataset and classification was conducted using all trials and leave-one-out cross-validation to test model performance. For each of the classifiers, the optimal cut-off value was defined as the point on the AUC where the product of sensitivity and specificity was maximized. We used *Shapley additive explanations* (SHAP) plots to explore the contributions of the different features to the predictions (Aas et al., 2021).We then repeated the classification, discarding the functional features related to frequency bands other than alpha and beta. The classifications carried out with the kNN were repeated using the SVM method (using a Gaussian kernel).

### Statistical analysis

The statistical significance of between-cohorts differences was examined at two levels, i.e., participant-wise (as in Fig. 1c) and parcel-wise (as in Fig. 1e). At either level, we used the non-parametric ANOVA (Kruskal-Wallis) test when the statistical comparison was conducted between three cohorts (NC, SCD, and MCI), and the Wilcoxon Rank-Sum test for post-hoc pairwise comparisons (NC-SCD, NC-MCI, SCD-MCI). To correct for multiple comparisons (false discovery rate, FDR), we used Benjamini-Hochberg method and estimated a threshold q=0.1 to calculate corrected p-values at participant-level and q=0.2 at parcel-level. True positives were then identified as corrected p-values less than significance level of 0.05. Similarly, the stats for correlation coefficients (Spearman or Pearson; mentioned in text, where employed) between DFA exponents and fE/I (e.g., in Fig. S3d) or between DFA exponent and structural atrophy/neuropsychological scores (Fig. 3a, b) were compared using paired t-tests, and FDR was used to account for multiple comparisons.

## Author contributions

SP, FM and JMP conceptualized the study. ISM and FM procured data. EJ, ISM and JVR preprocessed data. EJ analyzed data. GS carried out classification. EJ, JMP and SP conceived and prepared manuscript figures. EJ, ISM, GS, JMP, FM and SP wrote the manuscript. All authors approved the final manuscript.

## Data availability

The data analyzed during this study are stored at a secure UCM server accessible at the URL https://vbc.ucm.es/login.php. The access can be granted from the S. P. and F.M upon written request along with the clarification as to what purpose the data will be used.

## Code availability

The custom codes based on MATLAB (R2017b, Mathworks, Inc.), which are central to this Manuscripts’ results and conclusions are available as an open-source at https://github.com/palvalab/EI-unbalance-and-DFA-exponents.

## Extended data

**Extended data Fig. 1.**
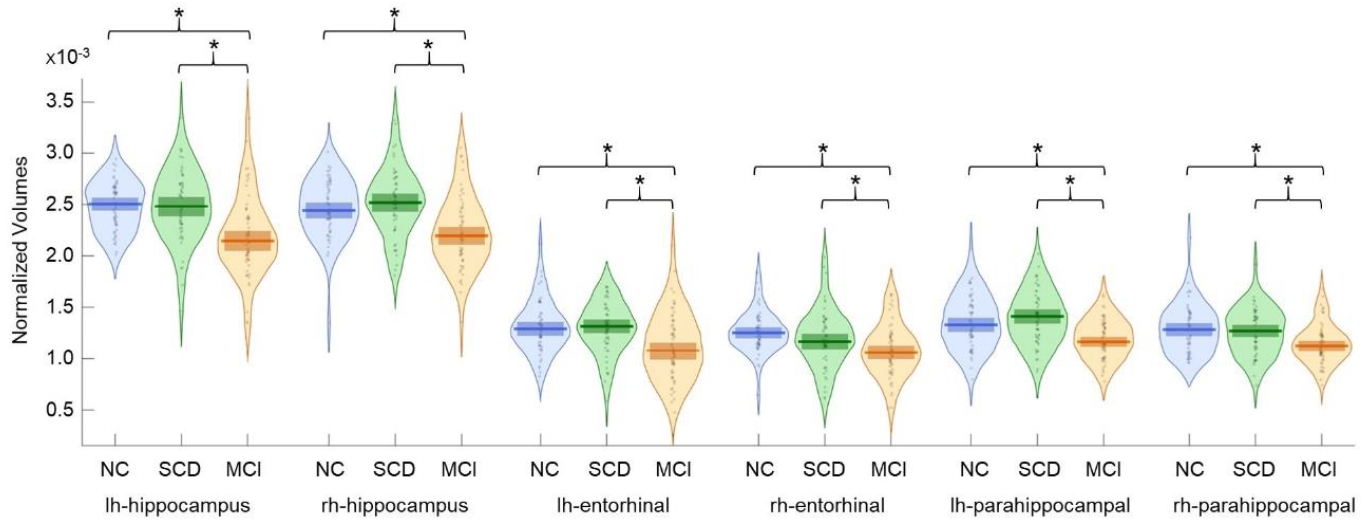
Structural atrophy across cohorts. Pairwise comparison shows reduced volume (* denotes p<0.05, Wilcoxon test) in MCI compared to NC and SCD cohorts in regions of the medial temporal lobe (hippocampal, entorhinal and parahippocampal). No significant differences were observed between NC and SCD.

**Extended data Fig. 2.**
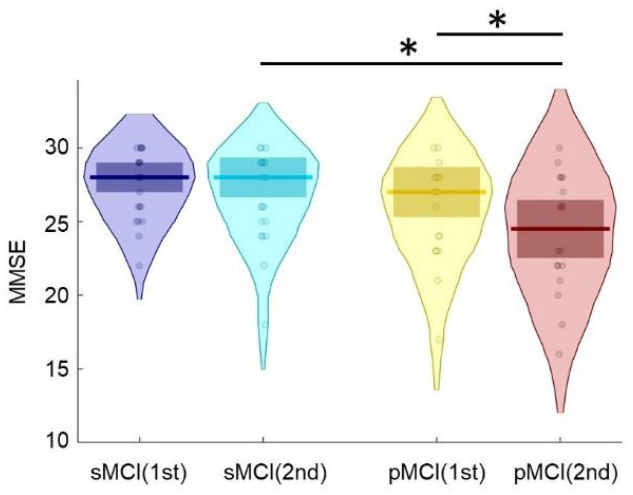
Decrease in MMSE score with disease progression. Differences between cohorts and measurements. * denotes significant (p<0.05, Wilcoxon test) differences between pairs.

**Extended data Table 1.**
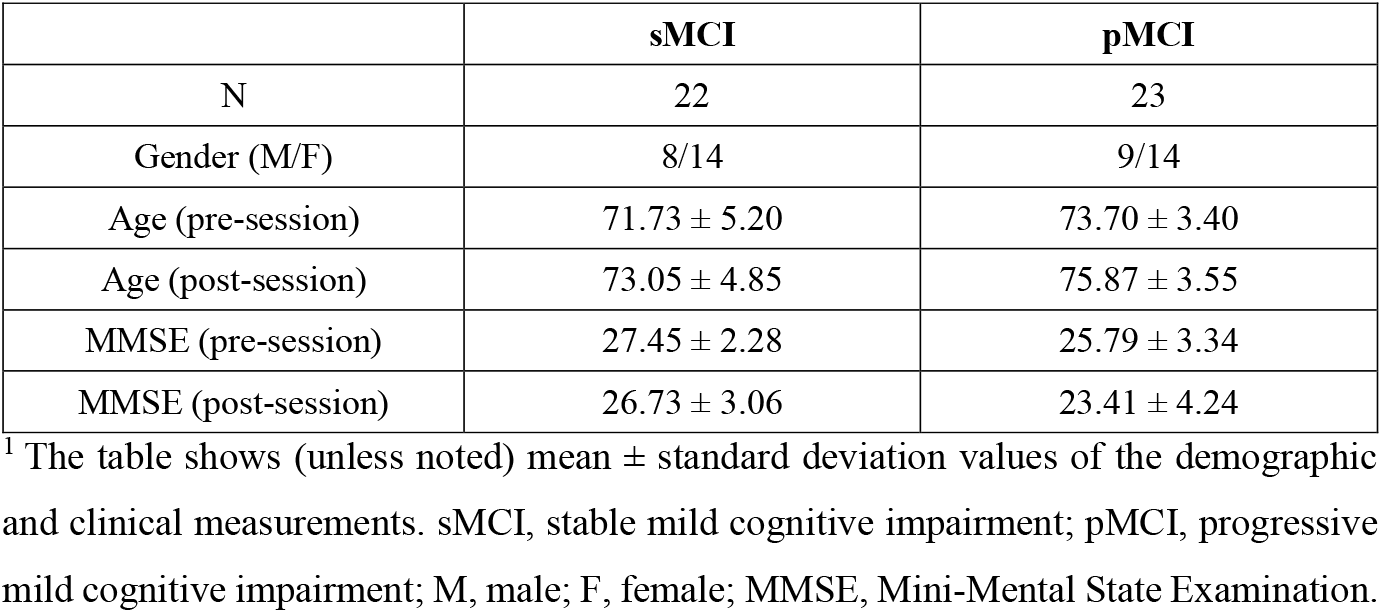
Demographic and clinical measurements for each cohort in the longitudinal dataset (pre- and post-sessions).

