## Supplemental Figures and Table for "Aberrant brain criticality as a neural basis of preclinical Alzheimer’s disease"

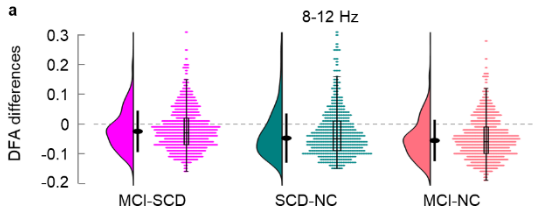


**Fig. S1| Between-cohorts DFA-differences distributions. a,** DFA exponent differences, pooled over individual frequencies in the 7-12 Hz range (where SCD differed from NC, as shown in (Fig. 1c)) as density plots (equally scaled heights across cohorts) where the black-filled dot denotes median, and the line length represents the standard deviation. Next to the density-plots are the observed individual subject's differences in DFA exponent (jittered) with an overlaid boxplot, where the box (from bottom-to-up) denotes the first quartile, the median and third quartile, while the whiskers length represents 1.5 times the interquartile range. **b,** Similar to (a), the plots show the between-cohort differences of DFA exponent for frequencies where SCD differed from MCI (12-22 Hz) whose averages are presented in (Fig. 1c).

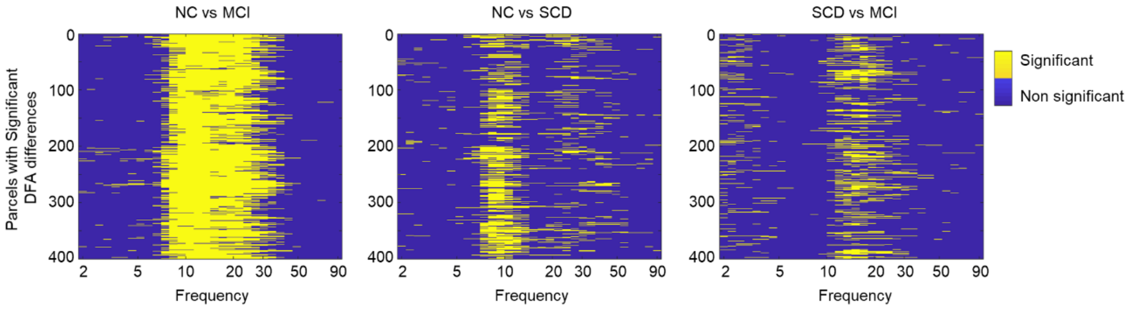


**Fig. S2| Parcel-level pair-wise DFA-exponents differences between cohorts.** Statistically significant ($p<0.05$) parcels-wise (Schaefer's 400) differences in DFA exponents between cohorts (left to right: NC vs MCI, NC vs SCD, and SCD vs MCI) are shown with yellow and non-significant with blue color. Wilcoxon rank-sum test was used for the statistical analysis for a given cohorts-pair and FDR was used for multiple comparison correction.

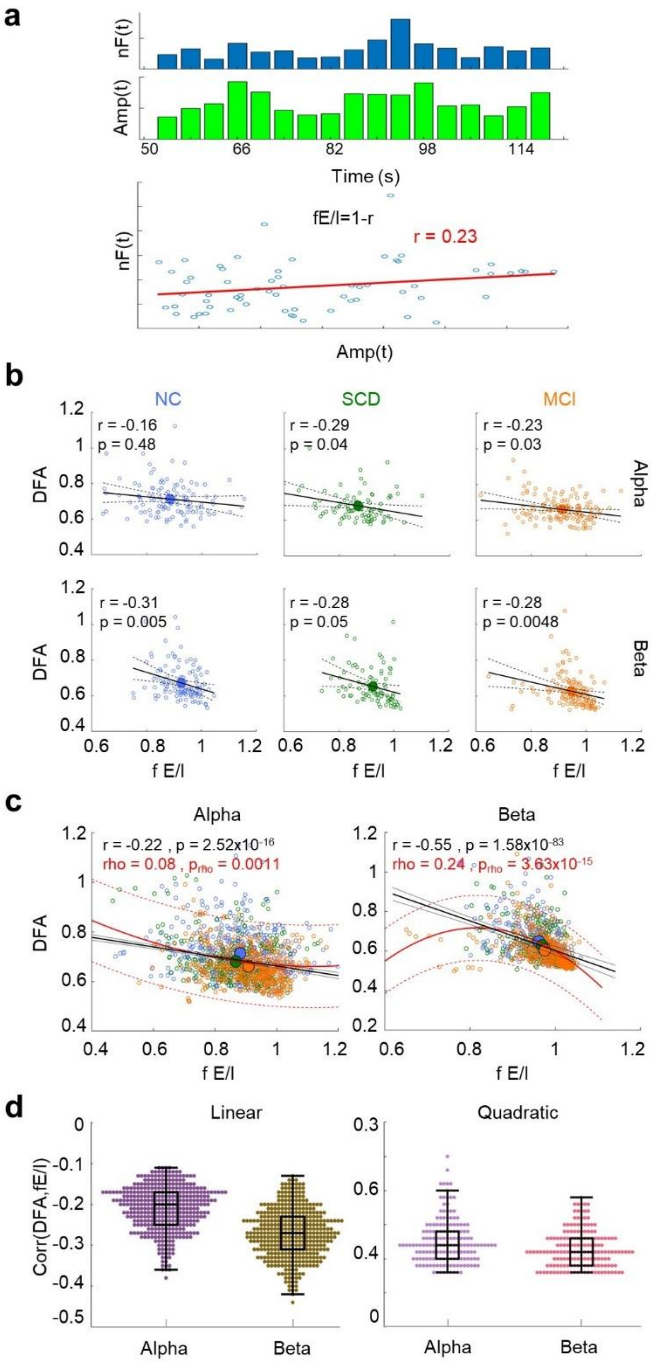


**Fig. S3| Dependence of DFA exponents on fE/I. a,** Normalized fluctuation function (nF(t)) and mean amplitude (Amp(t)) of each detrended signal profile, and Pearson correlation (r) between them, subtracted from 1 gives fE/I (see *Methods* for details). **b,** Negative linear correlation between individual groups' DFA exponent and fE/I. Solid black line represents association (Bonferroni corrected) along with dotted line (95% confidence interval) for the alpha and beta bands. **c,** Correlations for all individuals across cohorts. Mean DFA exponents and corresponding mean fE/I values of individuals for alpha and high beta bands. The larger filled circle represents the average across participants for each condition. rho denotes square root of goodness of fit (r2) for quadratic correlation, computed as the equivalent of linear correlation. **d**, Linear (left) and quadratic (right) correlations between DFA exponents and fE/I across participants. Dots representing significant ($p < 0.05$; FDR (Q=10%) corrected) association for given frequency.

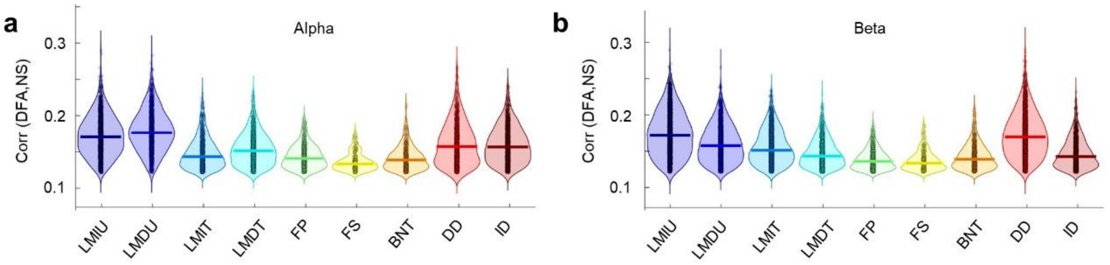


**Fig. S4| DFA exponents positively correlate with cognitive deficits.** Spearman Correlation between observed DFA exponents with neuropsychological scores (NS) after controlling for age confounds.

**a**

| Functional (F) | | | Structural (S) | Total features |
|---|---|---|---|---|
| Metrics | Networks | Bands | Volumes | F + S |
| 2 (DFA, fEI) | 7 (FPN, DMN, DAN, Lim, VAN, SM, Vis) | 4 (Theta, Alpha, Beta, Gamma) | 6 Left and right Hipp, Parahipp, Entrohinal | 62 |

**b** Features from alpha and beta bands: k-NN classifier and leave one out cross-validation

| Classes (n.trials) | Features (21) selected by mRMR (F=functional, S=structural) | Acc. | BAL acc | Pr. | Rec. | AUC |
|---|---|---|---|---|---|---|
| **NC-MCI** (116,142) | 15F + 6S | 0.78 | 0.78 | 0.81 | 0.77 | 0.84 |
| **NC-SCD** (116,85) | 15F + 6S | 0.78 | 0.78 | 0.74 | 0.75 | 0.79 |
| **SCD-MCI** (85,142) | 16F + 5S | *0.72* | *0.73* | *0.83* | *0.70* | 0.77 |

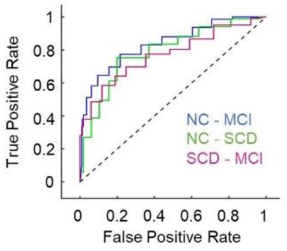


**c** Features from all bands: SVM classifier and leave one out cross-validation

| Classes (n.trials) | Features (15) selected by mRMR (F=functional, S=structural) | Acc. | BAL acc | Pr. | Rec. | AUC |
|---|---|---|---|---|---|---|
| **NC-MCI** (116,142) | 10F + 5S | 0.80 | 0.80 | 0.83 | 0.80 | 0.87 |
| **NC-SCD** (116,85) | 13F + 2S | 0.77 | 0.76 | 0.75 | 0.69 | 0.83 |
| **SCD-MCI** (85,142) | 12F + 3S | 0.79 | 0.80 | 0.89 | 0.76 | 0.87 |

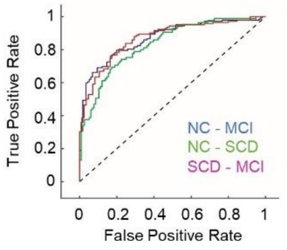


**d** Features from alpha and beta bands : SVM classifier and leave one out cross-validation

| Classes (n.trials) | Features (15) selected by mRMR (F=functional, S=structural) | Acc. | BAL acc | Pr. | Rec. | AUC |
|---|---|---|---|---|---|---|
| **NC-MCI** (116,142) | 9F + 6S | 0.76 | 0.77 | 0.88 | 0.66 | 0.82 |
| **NC-SCD** (116,85) | 12F + 3S | 0.72 | 0.71 | 0.68 | 0.64 | 0.75 |
| **SCD-MCI** (85,142) | 10F + 5S | *0.75* | *0.72* | *0.82* | *0.82* | 0.73 |

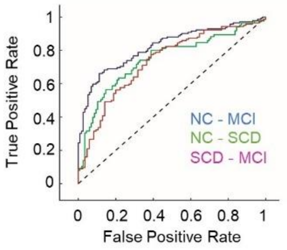


**Fig. S5| Consistency in classifier results using different set of features and classifiers.** **a,** Feature matrix: set of features subjected to the classifier for training and testing **b,** Classification results when features from alpha and beta bands (where most parcels were

significantly different) were classified using k-NN classifier and leave-one-out cross-validation along with ROC. **c,** The classification with the same features as in (Fig. 6c) but using support vector machine (SVM) classifier. **d,** SVM classifier's results with alpha and beta bands features. ROC curves for **(b)**, **(c)** and **(d)** are presented on their respective right.

**Table S1**. Neuropsychological battery tests description, ranges, and scoring criteria.

| Test | Score range [min, max] | Description |
|---|---|---|
| Texts of Verbal Memory (immediate units (LMIU), delayed units (LMDU), immediate thematic (LMIT), and delayed thematic (LMDT)) (Wechsler, 2004) | [ 0, 75 ] | Subjects are read 2 short texts and asked to pay attention since they will afterward be required to repeat the information with as much detail as possible. The second text is read twice. For the thematic scores (LMIT and LMDT) a checklist of 7 general ideas is used to score. Subjects get 1 point for each of the ideas that they are able to remember. The final score is a total of 7 points per text (7 × 3 in the LMIT since the second text is read twice, and 7 × 2 in LMDT). For the units scores (LMIU and LMDU) a checklist of 25 more specific ideas is used to score at 1 point each. The final score is 25 points per text (25 × 3 in the LMIU since the second text is read twice, and 25 × 2 in LMDU). The delayed scores were recorded 20 minutes after reading the texts. |
| Fluency Tests (phonemic (FP) and semantic (FS)) (Benton et al., 1983) | [ 0, — ] | In FP, subjects are asked to produce as many words as possible starting with the letters F, A, S (one minute for each). The primary score is the number of correct responses minus errors (i.e., repetitions or invalid responses such as proper nouns). The final score is the mean across the three letters. In FS, the letters were replaced by categories (*animals* and *fruits*). The final score is the mean of the two (one minute for each). |

| | | |
|---|---|---|
| Boston Naming Test (BNT) (Kaplan et al., 2001) | [ 0, — ] | Subjects are asked to name objects visually presented as two-dimensional line drawings in a booklet. The final score is the total number of identified objects within the initial 20 seconds. |
| Digits Span Test (forward (F) and backward (B)) (Wechsler, 2004) | F: [0, 16]<br>B: [ 0, 14 ] | Subjects are asked to repeat sequences of numbers in the same (F) and in reversed (B) order. Sequence lengths range from 2−9 digits for F and 2−8 digits for B, with two trials per sequence length. The final score is the number of correctly recalled sequences. |
